# Selection for later flowering time in an orchid through frost damage and pollinator activity

**DOI:** 10.1101/2024.02.14.580274

**Authors:** B.A. Wartmann, R. Bänziger, B. Györög-Kobi, K. Hess, J. Luder, E. Merz, B. Peter, M. Reutlinger, T. Richter, H. Senn, T. Ulrich, B. Waldeck, C. Wartmann, R. Wüest, W. Wüest, Q. Rusman, F.P. Schiestl

**Affiliations:** Arbeitsgruppe Einheimische Orchideen Aargau Schweiz (AGEO) c/o president Beat A. Wartmann, Sonnenbergstrasse 33, 8102 Oberengstringen; Department of Systematic and Evolutionary Botany, University of Zürich Zollikerstrasse 107. 8008 Zürich

**Keywords:** *Ophrys araneola*, *Andrena combinata*, selection, flowering time, pollination, frost damage, climate change

## Abstract

Flowering time is a key trait for plant reproductive success ensuring both overlap with pollinator activity and favorable conditions for fruit development. To quantify selection on flowering time, we individually marked 1250 plants of the Small Spider Orchid, *Ophrys araneola* RchB., in six populations in Northern Switzerland and surveyed them during three years. We recorded the date of first flowering, frost damage, and fruiting success of individual plants. In addition, we analyzed historical records of the orchid and its only verified pollinator, the solitary bee *Andrena combinata* in Northern Switzerland, to estimate potential desynchronization due to climate change. We documented strong selection for later flowering driven by frost damage, with all populations showing significant selection for later flowering in at least one year. Selection for later flowering driven by pollination (fruit set) could only be analyzed in one population due to the overall low fruit set, where it was significant in one year. The historical data from between 1970 and 2019 indicated low synchronization between orchid flowering and bee occurrence, with mean flowering three weeks earlier than the mean peak of bee occurrence, corroborating selection for later flowering through fruit set. The data also showed a significant advance of flowering time and bee-occurrence in the last decades, but to a similar degree in orchids and bees, hence without an indication of increasing desynchronization through climate change. Our study shows that selection for later flowering is mostly caused by frost damage, but also by the little synchronized flowering and pollinator activity, which is however unlikely to be a consequence of climate change in this orchid.

## Introduction

Flowering time is a key trait for plant reproductive success, and especially so for plants with specialized pollination system, as it must overlap with the emergence time of their pollinators, and favorable growing conditions to ensure successful seed maturation (Elzinga et al., 2007). Several studies have documented selection on flowering time, either in favor of earlier flowering (Internicola and Harder, 2012, Trunschke et al., 2017, Fogelstrom and Ehrlen, 2019, Agren et al., 2017), later flowering (O’Neill, 1997, Chen et al., 2017) or variable flowering time (Wu and Li, 2017, Parra-Tabla and Vargas, 2004). Sometimes, selection on flowering time has been linked to pollinator phenology or behavior; for example, selection for earlier flowering in food deceptive orchids is thought to be driven by flower-naïve pollinators, that have not yet learned to avoid unrewarding flowers (Internicola and Harder, 2012, Trunschke et al., 2017).

Overwhelming evidence shows that global warming impacts the phenology of plants and animals, with spring-flowering plants being particularly strongly affected (Fitter and Fitter, 2002, Molnar et al., 2012, Menzel et al., 2006, Cleland et al., 2007). These plants have advanced in flowering in the last decades by as much as 10 days (Robbirt et al., 2014, Bartomeus et al., 2011). Few studies up to now have investigated potential desynchronization between spring-flowering plants and their pollinators through climate change (Renner and Zohner, 2018). Whereas for plants with generalized pollination systems, little de-synchronization has been found (Forrest, 2015, Bartomeus et al., 2011, Forrest and Thomson, 2011, Razanajatovo et al., 2018), plants with more specialized pollination are more likely to suffer from increased mismatch between flowering and pollinator emergence. In the Japanese early-spring flowering, bumblebee-pollinated *Corydalis ambigua,* such a mismatch is generated when snowmelt is early but subsequent soil warming is slow, because bumblebee emergence depends on soil warming but flowering is triggered by surface warming after snowmelt. Therefore, plants flower earlier than bees emerge, and seed set is reduced (Kudo and Ida, 2013, Kudo and Cooper, 2019, Thomson, 2010, Kehrberger and Holzschuh, 2019, Robbirt et al., 2014). Another environmental factor affecting reproductive success in spring-flowering plants is frost damage of reproductive organs (Thomson, 2010). Such frost damage may increase with global warming, for example in alpine plants where earlier snowmelt may trigger earlier flowering and a higher exposure of buds and flowers to spring frost events (Inouye, 2008).

In principle, when organisms face low fitness because of changed environmental conditions, natural selection is expected to drive adaptive evolution towards a fitness peak, given traits under selection are heritable. Because variation in flowering time is controlled by genetic as well as environmental factors, selection on flowering time can lead to change in flowering time through adaptive evolution (Richardson et al., 2017, Anderson et al., 2012, Quinn and Wetherington, 2002). Whether plants can adapt to the rapidly changing environmental conditions caused by climate change is nevertheless little known, because few studies have quantified selection triggered by desynchronized flowering and pollinator activity or frost damage and the associated evolutionary response in plants (Thomson, 2010, Kehrberger and Holzschuh, 2019, Elzinga et al., 2007).

In our study, we investigated the Swiss native orchid, *Ophrys araneola* RchB. and its only verified pollinator, the solitary bee *Andrena combinata* (Christ 1791) (Schiestl and Vereecken, 2008). Like almost all species of the genus *Ophrys, O. araneola* is fully outcrossing and employs sexual mimicry for pollination, by imitating the mating signals of female bees and thus attracting males that attempt to copulate with the flower labellum (Johnson and Schiestl, 2016). Pollination by sexual mimicry is often species-specific, and floral scent has been shown to be a key trait for pollinator attraction, with often species-specific composition, fine-tuned for the attraction of one or a few pollinator species (Ayasse et al., 2011, Schiestl, 2005, Dötterl and Vereecken, 2010). To investigate selection on flowering time we conducted surveys in six natural populations in Switzerland, documenting flowering time, frost damage and fruiting success in 1250 individually marked plants over three years in the field. To study potential effects of climate change on desynchronization between plant flowering and pollinator activity, we analyzed historical occurrence data of the orchid and its pollinator bee in the region of the study area. We addressed the following specific questions in our study: 1) Is the survival of inflorescences and fruiting success in this orchid associated to the time of flowering, leading to selection on flowering time? 2) Did the flowering time of *Ophrys araneola* and the emergence time of *Andrena combinata* change in the last decades in Switzerland, leading to increased desynchronization between orchid and its pollinator?

## Materials and Methods

### Flowering time and temperature in contemporary populations

In February 2021, between 100 and 300 rosettes (1250 in total) of *O. araneola* in 6 populations in the Swiss cantons of Aargau and Zürich (Table 1), where possible close to existing tracks in the field, were marked individually with metal tags. In some populations, newly discovered plants were marked in 2022 and 2023. Because not every plant flowers every year, and many plants got damaged, especially by frost events, and could not complete flowering, the total number of records obtained from plants throughout the three study years was considerably less than 3 × 1250. One data logger was placed in each population, close to where *O. araneola* plants were growing, to record hourly temperatures. To avoid losing loggers when populations were mowed during late summer/autumn, data logger were removed after flowering, and placed again into the field at the end of the year. All metal tags and data loggers were removed from the field in summer/autumn 2023.

**Table 1.** *Ophrys araneola* populations surveyed in this study. All populations are found on southerly facing slopes, with an altitude of 370 to 540 m above sea level, calcareous soil and semi-dry meadow plant communities; the populations are annually mowed late in September allowing maturation of orchid seeds.

| Population | Swiss grid, X-coordinate | Swiss grid, Y-coordinate | Latitude | Longitude | Altitude |
| --- | --- | --- | --- | --- | --- |
| Erlinsbach (Lehrpfad) | 2642920 | 1251590 | 47° 24' 49.40" | 8° 0' 26.39" | 530 m |
| Kloten (Feek, Eigental) | 2689260 | 1258640 | 47° 28' 20.91" | 8° 37' 21.71" | 540 m |
| Birmenstorf (Schluu) | 2661610 | 1257460 | 47° 27' 54.12" | 8° 15' 20.79" | 460 m |
| Küttigen (Judenhalde) | 2646320 | 1252660 | 47° 25' 23.22" | 8° 3' 8.97" | 470 m |
| Villigen (Cheestel) | 2658810 | 1265670 | 47° 32' 20.85" | 8° 13' 11.01" | 370 m |
| Villnachern (Bahnbord) | 2654420 | 1258420 | 47° 28' 27.48" | 8° 9' 37.93" | 410 m |

### Flowering time and fruiting success

Subsequently, for three years, surveys of flowering time and fruiting success in the populations were conducted. Surveys were done every 2-3 days from early March to end of May, where start of flowering of marked plants was recorded. Plants without inflorescence in any particular year were noted as not flowering. Frost damage was assessed during the surveys by recording plants with wilted inflorescences that showed no signs of snail- or mechanical damage (Figure S1). When frost only partially damaged an inflorescence and the plant continued to flower, this was not counted as frost damage. Thus, our record of frost damage included only plants where the inflorescence had been fully destroyed and subsequently dried out without producing any more flowers in the respective flowering season. Such damage obviously prevented the plant from producing any fruits in the particular season but did not necessarily impact re-flowering in the next season (see results section). After the flowering period (i.e. by mid-May-June), total number of flowers (wilted, dried flowers are still visible after flowering), number of fruits, and plant height was recorded from all plants that had produced an inflorescence and had not been damaged by frost.

To confirm the presence and identity of pollinators in the populations, we attempted to catch pollinators in all populations by using hand nets and by searching for bees carrying pollinaria on the flowers and in the surrounding of flowering *O. araneola* plants. All in all, three male bees were caught in the Birmenstorf population, with two of them carrying pollinaria of bright yellow color typical for the genus *Ophrys.* The bees were identified using a reference specimen of *A. combinata* and the key provided in (Amiet et al., 2010). One pollinarium was sequenced using ITS2 primers with standard PCR and sanger sequencing methods (Breitkopf et al., 2015). The sequence was compared to the DNA extracted from an *O. araneola* leaf.

### Historical orchid/bee records

Historical records of *Ophrys araneola* in Switzerland were obtained from the database “InfoFlora” (www.infoflora.ch). These included records from herbaria dating back to 1868, as well as more recent records collected by professionals and plant enthusiasts in the field. *O. araneola* is sometimes considered a subspecies of *O. sphegodes*, and historically (i.e. before 1950 (Schmid, 1998)) has not always been treated as a distinct species. Therefore, our historical data may contain some *O. sphegodes* records, but as *O. sphegodes* is much rarer than *O. araneola* in the area considered for our study (www.infoflora.ch), the historical dataset of *O. araneola* can be considered reasonably accurate. Records of the pollinator bee, *Andrena combinata* (males and females) were obtained from the Swiss database “InfoSpecies” (Swiss National Apoidea Databank https://doi.org/10.15468/ksfmzj); as for the orchid data, these data comprise specimen of insect collections and records from bee-collectors, going back to the 1970s. The data did not indicate the gender of the bees, making a male-specific analysis impossible; although in several bee species, males emerge before females, this seems to be not be the case for *Andrena combinata* (Westrich, 1989, Schiestl and Vereecken, 2008), making a combined analysis of males and females reasonable. For the analysis of flowering advancement, all orchid data were used; to analyze desynchronization, only the time period for which orchid and bee data were available (i.e. from 1970 to 2019) was used; in addition, to achieve a sample from a climatically and topographically homogeneous region, we only used data from Northern Switzerland (comprising the greater region where the field work was conducted, excluding the north-western region,). Specifically, 52 bee records and 725 orchid records from the cantons of St. Gallen, Thurgau, Schaffhausen, Zürich, Aargau, Luzern, Schwyz, Solothurn, Basel-Land, and Bern were used.

## Statistical analysis

### Data of population surveys

Generally for flowering time/bee emergence, all date-values were converted into “day of the year” to allow statistical analysis of flowering time as an ordinal variable. For the contemporary data, temperature data obtained from the data logger were used to calculate mean temperatures for the months January – June per population throughout the three study years. For the rest of the months, not enough data were available to allow a reasonable analysis. For the mean temperature of each month, a maximum of 18 data points were created (6 populations and 3 years). However, because data loggers were lost in the field or ran out of battery power, the full 18 data points were not available for any months. Pearson product moment correlations were then calculated between mean temperatures of the 6 months that were analyzed and the mean “start of flowering” and “earliest flowering time” in the populations. Pearson product moment correlations were also calculated for the measured traits of individuals between pairs of years (i.e. “start of flowering”, “number of flowers”, “plant size”). Spearman rank correlations were used for binary (frost damage, flowering) and count variables (number of fruits).

### Selection via frost damage

Factors impacting frost damage were assessed by a generalized linear mixed model (GLMM), with frost damage as the (binary) response variable (no-damage/damage). “Population” and “year” were included as factor, “day of first flower” as covariate, and the interaction between “day of first flower” and “population”, and “day of first flower” and “year”. Selection coefficients on flowering time via survival of inflorescences through frost were estimated as using binary logistic regression, for each population and year separately, to allow for the estimation of population- and year-specific selection. In total, 16 selection coefficients (out of 18 year/population combinations) were calculated, because in two populations, no frost damage was recorded in 2023.

### Selection via fruiting

**success** Because few plants had fruits, and the number of fruits was generally low (mean (±s.e.m) = 0.07 (±0.10); see Table S1), the numerical fruit set data was transformed into a binary data (no-fruits/fruits), to avoid problems with many zero values in the analysis. Factors impacting fruit-set were assessed by a generalized linear mixed model (GLMM) with binary distribution with “no-fruits/fruits” as dependent variable. Plants with frost damage were not included in this analysis, as they did not successfully complete flowering. “Population” and “year” were included as factors, “day of first flowering”, “number of flowers” and “plant height” as covariates; the interaction between “day of first flowering” and “population”, and “day of first flowering” and “year” was also included in the model. Selection coefficients on flowering time via reproduction were estimated by binary logistic regression. This was only done for the population “Birmenstorf”, as for the other populations, the number of plants with fruits was considered too low for a meaningful analysis (see Table S1). In the selection analysis, we used fruits rather than seeds as fitness variable, as seeds cannot easily be counted in orchids, that typically produce tiny dust-like seeds by the thousands. Relative fitness (i.e. fitness of an individual divided by the mean fitness of the population) was also not considered in our study, because the mean number of fruits was not significantly different between populations across all years (F_1,1526_ =1.21, P=0.271), and the calculation of relative fruit set would have led to very high values in few plants, as the number of fruits was low with many plants having no fruits in all populations (see Table S1), leading to unusual data distributions. Orchids pollinated through deceptive pollination mechanisms, like all members of the genus *Ophrys*, often have very low fruit-set and are highly pollen-limited in their reproduction (Scopece et al., 2010, Scopece et al., 2015, Johnson and Schiestl, 2016). Therefore, hand-pollination experiments to assess the degree of pollen-limitation in our study populations were not performed.

### Historical data

To assess the flowering time of *O. araneola* throughout the time period of the historical records, the data was sorted according to its year into four time-categories, A: 1868-1900, B: 1901-1940, C: 1941-1980, D: 1981-2019, and the mean values for the dates of those records was calculated. To estimate any change in flowering time throughout the last 150 years, a linear regression with “day of the record” as dependent variable and “year of record” as independent variable was calculated (a quadratic regression was also done and led to very similar values). In this analysis, a negative slope of the regression line (β) indicates an advance in the dates of the records, and a positive slope indicates a delay in the dates. To compare historical bee- and orchid data, only data from the same time period (1970-present) and geographic region were used (north-eastern Switzerland, the region in which the study was conducted). To analyze difference in the change of orchid-flowering and bee-occurrence, a general linear model was used with “day of the record” as dependent variable, “species” (i.e. orchid/bee) as fixed factor, “year” as covariate, and the interaction between “species” and “year”. Here, a significant interaction between “species” and “year” indicates a difference in the way “date of the record” and “year” are associated with each other in orchids and bees. This would indicate a difference in the way orchid flowering and bee emergence changes throughout time. To compare flowering time and bee emergence, a one-way ANOVA with “date of the record” as dependent and “species” as factor was conducted. All statistical analyses were performed with IBM SPSS statistics version 14.

## Results

### Data of population surveys

In our study populations, mean start of flowering varied considerably, from 25^th^ March (day 84) in Villnachern in 2022, to 25^th^ April (day 115) in Kloten in 2021 (Table 2, Table S1). Start of flowering of the same individuals was correlated highly significantly among the years: 2021-2022: r_372_=0.84, 2021-2023: r_342_=0.71, 2022-2023: r_402_=0.67, all P<0.001). The same was found for “number of flowers” and “plant size”, where values of individuals between years were highly significantly correlated (number of flowers, 2021-2022: r_224_=0.38; 2021-2023: r_237_=0.40, 2022-2023: r_302_=0.39 all P<0.001; plant size, 2021-2022: r_120_=0.46, 2021-2023: r_101_=0.40, 2022.2023: r_233_=0.40, all P<0.001), indicating constant environments and/or heritability of these traits. Mean temperature in February showed the strongest correlation with mean start of flowering, suggesting later flowering with lower temperatures in February (Table 2). This association was also highly significant for “earliest flowering”, suggesting temperatures in February impact the time of flowering in this orchid species.

**Table 2.** Correlation between flowering time and average temperatures in January-June in the six populations, assessed within three years. Values show Pearson product-moment correlation coefficients (r), with p-values in brackets. Significant values are shown in bold. Numbers behind months are number of data points used in the analysis.

| Mean temperature per month (no. data points) | Mean start of flowering | Earliest flowering |
| --- | --- | --- |
| January (9) | -0.48 (0.192) | -0.25 (0.518) |
| February (10) | **-0.70** (0.024) | **-0.656** (0.040) |
| March (13) | -0.03 (0.930) | 0.07 (0.815) |
| April (16) | -0.42 (0.105) | -0.40 (0.125) |
| May (16) | **-0.50** (0.046) | **-0.52** (0.040) |
| June (10) | -0.21 (0.570) | -0.51 (0.133) |

### Frost damage

Frost damage was variable and reached devastating levels with over 80% of plants damaged and unable to complete flowering in some populations and years (Figure 1). Frost damage of individuals was correlated among years (2021-2022: ρ_400_=0.37; 2021-2023: ρ_349_=0.29; 2022-2023: ρ_352_=0.19, all P<0.001), indicating many individuals were damaged in multiple years. Frost damage in 2021 had no effect on flowering of individual plants in 2022 (ρ_541_=-0.05, P=0.210), but frost damage in 2022 had a significant negative effect on the likelihood of flowering in 2023 (ρ_510_=-0.13, P=0.005).

**Figure 1.**
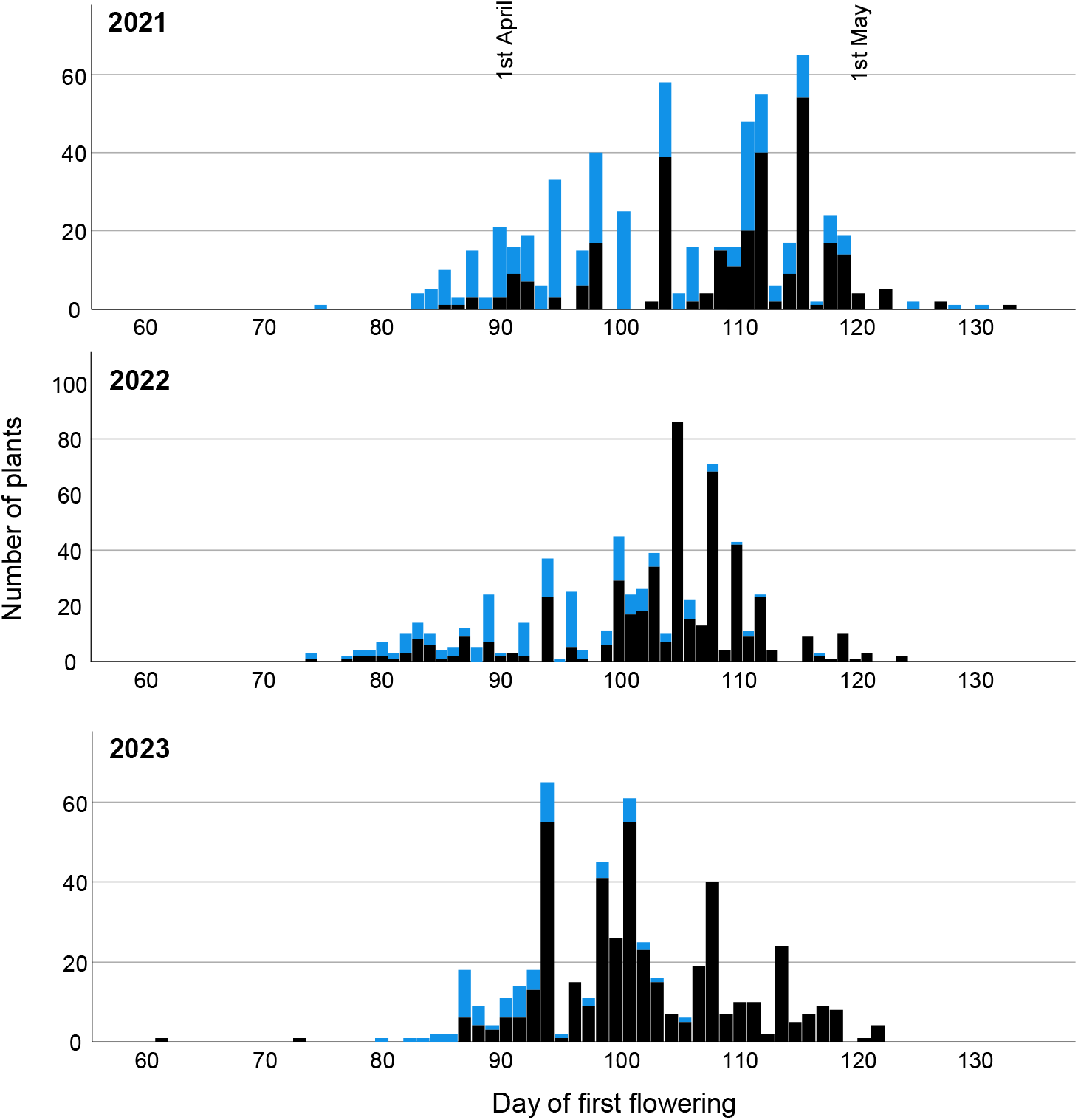
Association between flowering time and frost damage in plants in all six study populations, shown separately for the three study years. Plants suffering frost damage are indicated as the top, light blue section of the bars. The figure shows that frost damaged decreased with later flowering throughout all three years, albeit with different levels of significance (for statistical values see Tables 3 and 4).

**Table 3.** Factors impacting frost damage in the study populations assessed by a generalized linear mixed model with binary distribution. Frost damage (no-damage/damage) was used as dependent variable, “population” and “year” as factors, and “day of first flower” as covariate. Interactions between factors and the covariate were included, too. Besides population and year, flowering time had a highly significant impact on frost damage, with early flowering plants being more affected than later flowering ones (see also Figure 3). A total of 1978 plant records, based on 1147 individually marked plants were included in this analysis (some plants were assessed during more than one year).

| Source | Chi<sup>2</sup> | df | P |
| --- | --- | --- | --- |
| **Population** | **11.55** | **5** | **0.041** |
| **Year** | **17.08** | **2** | **<0.001** |
| **Day of first flower** | **114.17** | **1** | **<0.001** |
| Population x Day of first flower | 7.61 | 5 | 0.179 |
| **Year x Day of first flower** | **25.64** | **2** | **<0.001** |

**Table 4.** Estimated selection coefficients (± s.e.) on flowering time caused by frost damage for all populations and years separately. Values in brackets are number of plants included in the analysis for each population. Two populations in 2023 (Erlinsbach, Birmenstorf) had no plants with frost damage and were thus not included in this analysis. For the binary logistic regression analysis, frost damage (no-damage/damage) was used as dependent variable and “day of first flower” as independent variable. Significant coefficients are shown in bold.

| Year | Population | Chi<sup>2</sup> | df | coefficient | P |
| --- | --- | --- | --- | --- | --- |
| 2021 | Erlinsbach (51) | 7.44 | 1 | **-0.14±0.05** | **0.006** |
| | Kloten (135) | 0.01 | 1 | 0.00±0.06 | 0.946 |
|  | Küttingen (65) | 6.04 | 1 | **-0.09±0.03** | **0.014** |
| | Birmenstorf (102) | 2.51 | 1 | -0.13±0.08 | 0.113 |
|  | Villigen (158) | 19.97 | 1 | **-0.22±0.05** | **0.000** |
|  | Villnachern (73) | 10.95 | 1 | **-0.18±0.05** | **0.001** |
| 2022 | Erlinsbach (50) | 2.23 | 1 | -0.13±0.08 | 0.135 |
|  | Kloten (215) | 1.90 | 1 | -0.25±0.18 | 0.169 |
|  | Küttingen (72) | 9.76 | 1 | **-0.15±0.05** | **0.002** |
|  | Birmenstorf (61) | 5.133 | 1 | **-0.13±0.06** | **0.023** |
|  | Villigen (190) | 38.46 | 1 | **-0.17±0.03** | **0.000** |
|  | Villnachern (68) | 6.66 | 1 | **-0.15±0.06** | **0.010** |
| 2023 | Kloten (52) | 5.55 | 1 | **-0.46±0.19** | **0.018** |
|  | Küttingen (85) | 13.73 | 1 | **-0.35±0.09** | **0.000** |
|  | Villigen (172) | 14.80 | 1 | **-0.22±0.06** | **0.000** |
|  | Villnachern (70) | 5.51 | 1 | **-0.12±0.05** | **0.019** |

Our analysis showed that “day of first flower” (i.e. flowering time) highly significantly explained frost damage; as expected, early flowering individuals were much more likely to be damaged by frost (Table 3; Figure 1). “Population”, “year”, and the interaction between “year” x “day of first flower” were also significant in the analysis, suggesting frost damage varied especially among years and the impact of flowering time on frost damage varied among years. In the analysis with years and populations separately, 12 out of 16 year/population combinations showed a significant association between flowering time and frost damage (Table 4; 2 populations in 2023 had no frost damage). The estimated selection coefficients varied from 0.46 to 0.09.

### Fruiting success

Fruit set was low in most populations (Table S1); the percent of individuals with at least one fruit ranged between 15 (Birmenstorf in 2021) to 0 (plants in Erlinsbach did not have any fruit set throughout all study years, except one plant which set 2 fruits in 2022, and two plants that set a total of three fruits in 2023, both outside the study area), with a total mean (± s.d.) of 3.43 (±4.58) for all populations and years (Table S1). The mean number of fruits produced by all plants of a population varied between 0.32 (±0.73) (Birmenstorf) and 0, with a total mean of 0.07 (±0.10) for all populations and years. Number of fruits produced by individuals was significantly positively correlated between 2021 and 2023 (r_238_=0.31, P<0.001), and between 2022 and 2023 (r_318_=0.15, P=0.008). Nevertheless, the number of fruits produced in one year had a significant negative effect on the likelihood of flowering in some of the following years (2021-2022: ρ_337_=-0.14, P=0.012; 2021-2023: ρ_389_=-0.05 P=0.390; 2022-2023: ρ_362_=-0.18, P<0.001; Figure 2), and on number of flowers produced in some of the following years (2021-2022: ρ_217_=-0.18, P=0.009; 2021-2023: ρ_232_=-0.17, P=0.008; 2022-2023: ρ_301_=0.02, P=0.680), suggesting a trade-off between fruiting and plant performance in the following years.

**Figure 2.**
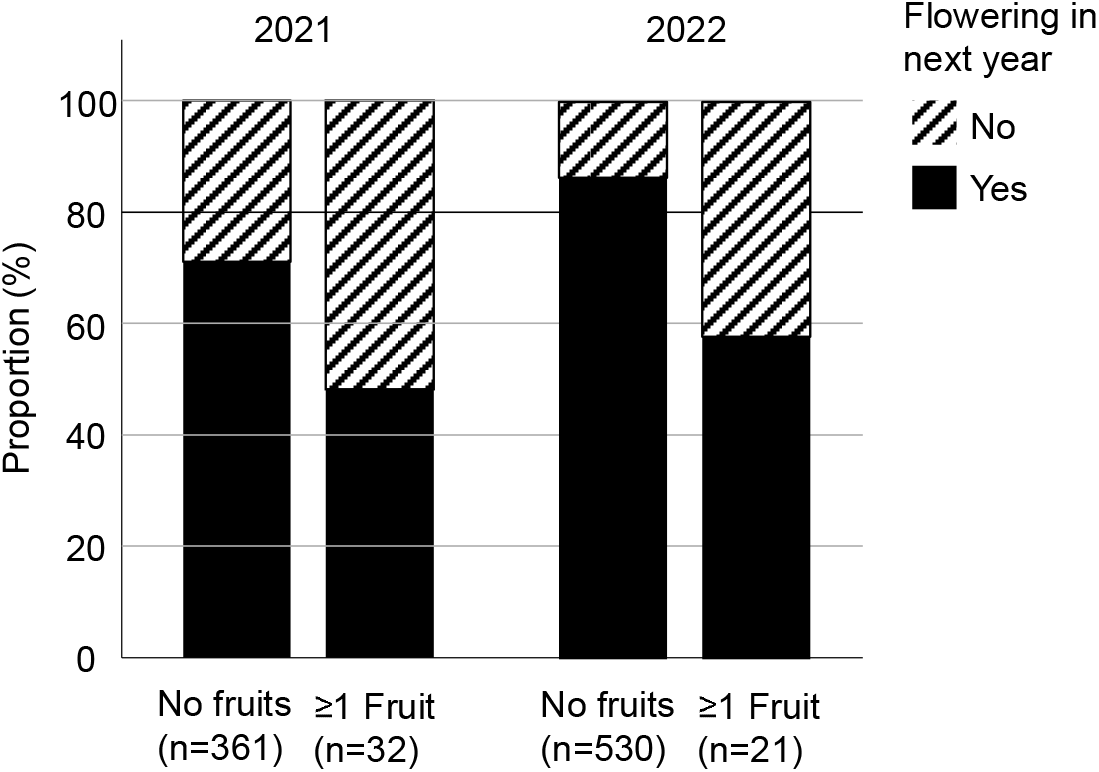
Plants that produced one or more fruits had a reduced likelihood of flowering the next year, indicating a trade-off between fruiting and subsequent flowering (see results section for statistical values).

The likelihood of producing at least one fruit was different between populations and years, and significantly associated with plant size (Table 5). Whereas flowering time was not significant in this analysis including all populations, the interactions between flowering time and year, and flowering time and population were significant (Table 5). When the analysis was done for the population with the highest fruit set only (Birmenstorf), the likelihood of producing at least one fruit showed a significant association with flowering time in 2021, with a higher likelihood of producing fruits for later flowering individuals (Figure 3; Table 6).

**Figure 3.**
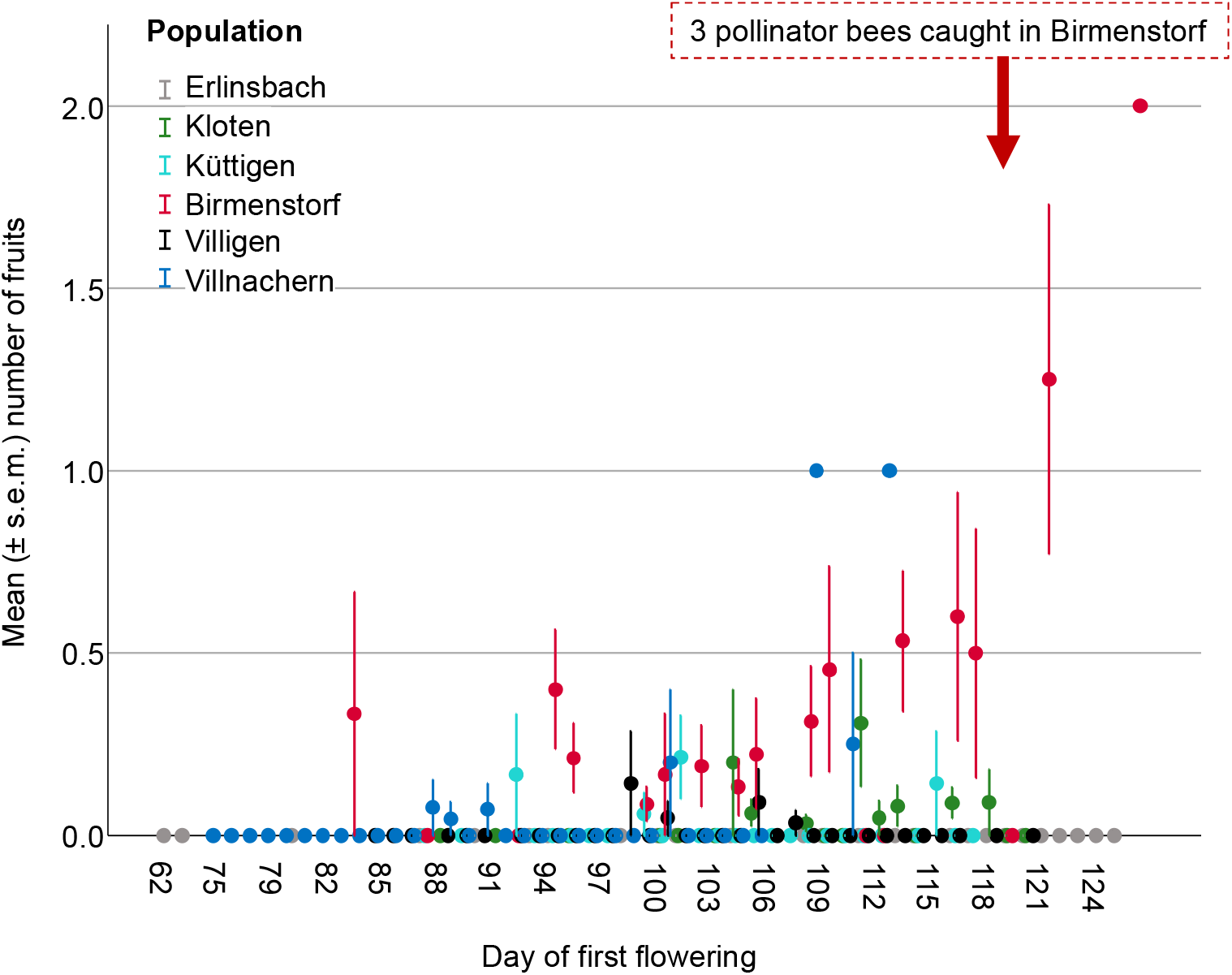
Association between mean number of fruits produced by plants in the study populations and their “day of first flowering”. Fruiting success differed between year and population, and was positively associated to plant size, and “day of first flower” in a population- and year-specific way (significant interaction between “days of first flower” and “year”, and “day of first flower” and “population”; for statistical values, see Table 5 and Table 6).

**Table 5.**
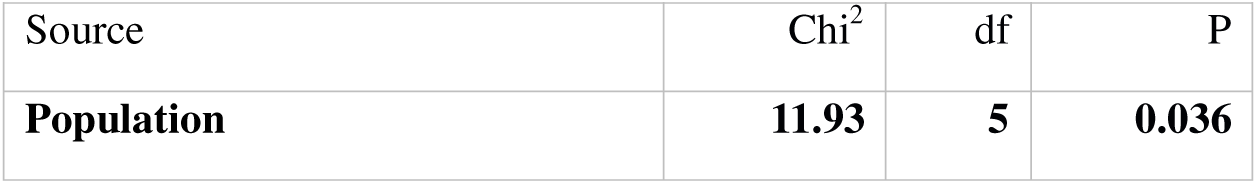
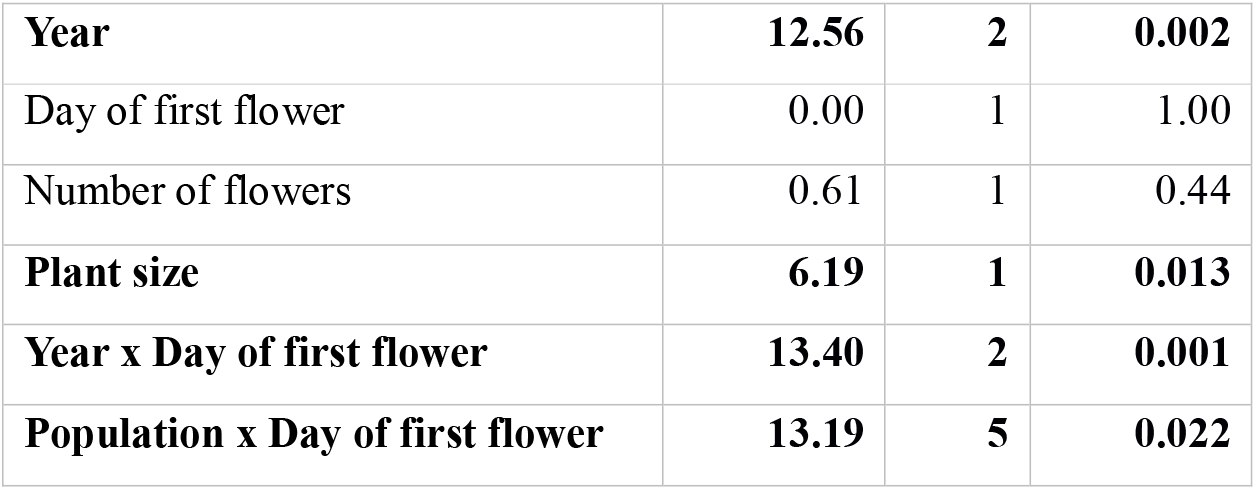
Factors impacting the number of fruits produced by the plants assessed by a generalized linear mixed model with binary distribution (no-fruits/fruits), using plants of all populations. “No-fruits/fruits” was used as dependent variable, “population” and “year” as factors, and “number of flowers”, “plant size” and “day of first flower” as covariates. Significant factors are shown in bold. This analysis was based on a total of 987 individually marked plants (Erlinsbach: 90, Kloten: 330, Küttigen: 114, Birmenstorf: 168, Villigen: 208, Villnachern: 77) that produced a total of 1528 inflorescences that completed flowering without frost damage throughout the three study years.

**Table 6.** Estimated selection coefficients (± s.e.) on flowering time through fruit production in the population Birmenstorf. Values in brackets are number of plants included in the analysis for each year. For logistic binary regression, fruit production (no-fruits/fruits) was used as dependent-, and “day of first flower” as independent variable. Significant coefficients are shown in bold.

| Year | Chi2 | df | coefficient | P |
| --- | --- | --- | --- | --- |
| 2021 (102) | 15.71 | 1 | **0.19±0.05** | **<0.001** |
| 2022 (64) | 2.81 | 1 | -0.08±0.05 | 0.094 |
| 2023 (82) | 1.23 | 1 | 0.05±0.04 | 0.263 |

Three males of *Andrena combinata* were caught on 29^th^ April 2023 in the population Birmenstorf (Figure 4). Two of the bees carried pollinaria, the ITS2 sequence of which were identical to the one derived from an *O. araneola* leaf (Figure S2). ITS2 is not species-specific in the genus *Ophrys* but differs between *O. araneola* and *O. insectifera* (Bateman et al., 2003), with *O. insectifera* being the only co-flowering *Ophrys* species during the time of the study in the Birmenstorf population.

**Figure 4.**
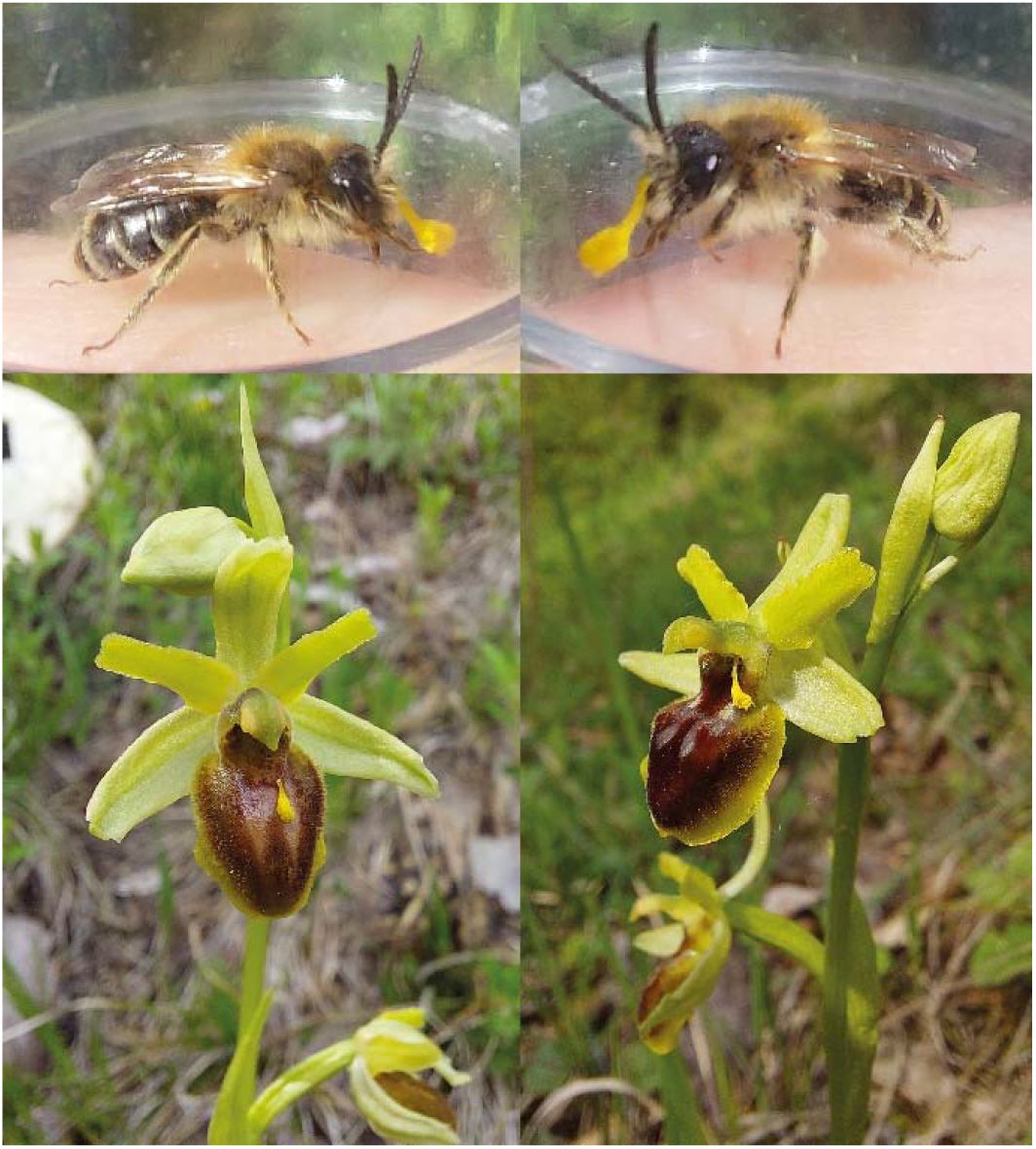
Pictures of one individual male of *Andrena combinata* carrying pollinaria of *Ophrys araneola*. The identity of the pollinaria was confirmed by ITS2 sequences. Below are two flowers of *Ophrys araneola* with one loose pollinarium, apparently deposited by the pollinator bees. Pictures of both bees and orchids were taken on April 29^th^ 2023 in the population Birmenstorf.

### Historical data: Flowering time and activity of the pollinator bee

Our analysis of historical *O. araneola* records showed that within the last 150 years, the flowering time of this orchid has advanced by about 12 days in Switzerland (Figure 5A shows only the last 50 years, i.e. the time where bee records were available, too). The mean “time of the record” in four year-categories changed from 12^th^ May in the 19^th^ century (1869-1900; day 132±10.42, n=16) to 8^th^ May (1901-1940; day 128±10.2, n=32, and 1941-1980; day 128±16.23, n=164) to 30^th^ April (1981-2019; day 120±17.58, n=836). Using data of all individual years, a highly significant advance in flowering time was detected (β=-0.24, P<0.001). The analysis for the time period where both orchid and pollinator-bee data were available (1970-present) showed a similar advance of flowering time in orchids and emergence time in bees (bee: β=-0.31, P=0.030, orchid: β=-0.38, P<0.001; Figure 5A). There was no significant difference in the way flowering time and bee-emergence advanced during this time-period (interaction species and year: F_1_=0.25, P=0.620). Comparing the abundances of records throughout the flowering/bee-activity season showed that orchids flowered consistently earlier than bees emerged, with the earlier flowering orchids well ahead of the first bees (earliest orchid record: 5^th^ March (3^rd^ March in our data), earliest bee record: 16^th^ April (Figure 5B). The mean flowering time of the orchid was also significantly earlier than emergence time of the bee (F_1,717_=77.8, P<0.001).

**Figure 5.**
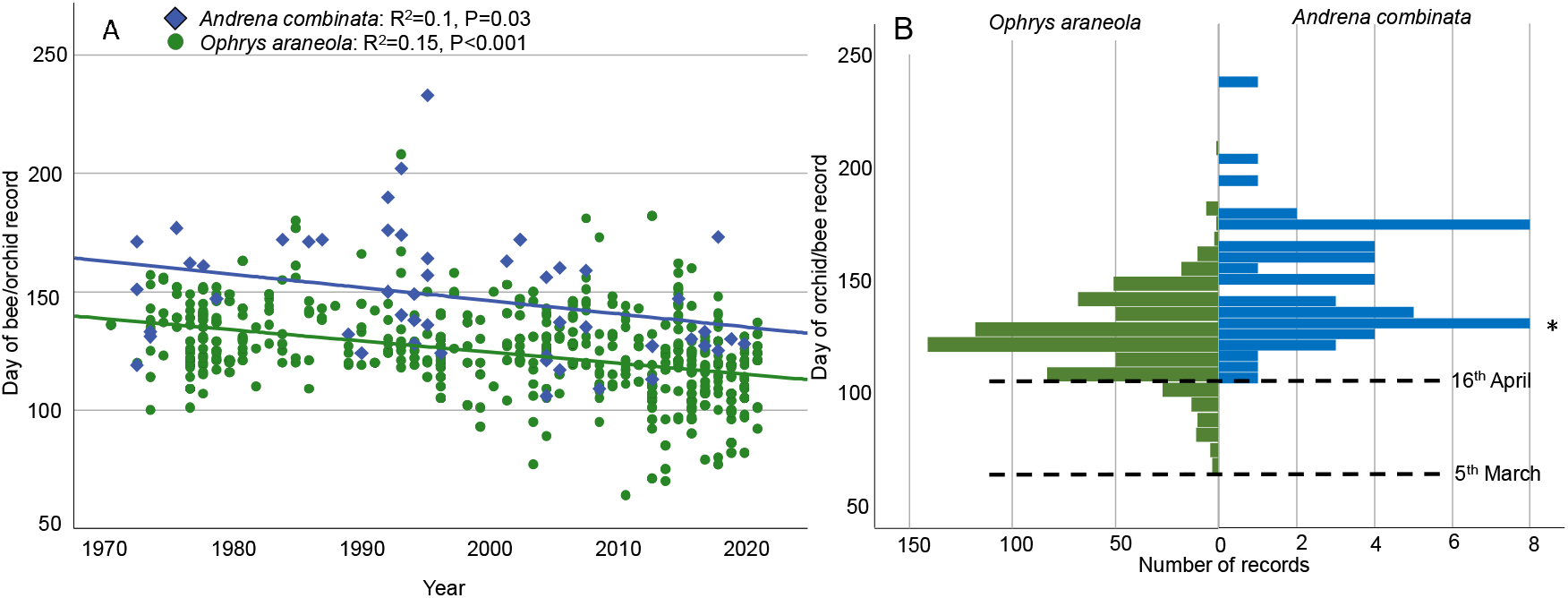
Historical records of *Ophrys araneola* and its pollinator, *Andrena combinata* in Northern Switzerland (data obtained from www.infoflora.ch, www.infospecies.ch). A) Date of record drawn against the year of the record, showing that both, flowering time of the orchid and the emergence of the bee, have advanced similarly in the last 50 years in Switzerland. B) Histogram of abundances of records at the different dates, showing the orchid to flower significantly earlier (ca 20 days) than the bees to emerge (mean±s.d., orchid: 123.64±18.28 (3^rd^ May), bee: 147.63±25.46 (27^th^ May); F_1,717_=77.8, P<0.001). * on 29^th^ April 2023, three males of *Andrena combinata* were caught at the population Birmenstorf, two of which carried pollinaria of *O. araneola*.

## Discussion

Natural selection through reproduction and survival has been commonly documented in plants and is thought to be fueled by biotic factors such as pollinators or herbivores, as well as abiotic factors such as climate (Caruso et al., 2019, Trunschke et al., 2017, Agren et al., 2017, Fogelstrom and Ehrlen, 2019). Whether organisms can respond to selection through adaptive evolution depends on the strengths of selection and the evolvability of a trait, which is its genetic variance and covariance (Conner and Hartl, 2004) versus the strengths of environmentally-induced plasticity (van Kleunen and Fischer, 2005). Our unique dataset is based on surveys of individually marked orchids over three years, and documents selection for later flowering through both survival of inflorescences (escape of frost damage) and reproduction (i.e. fruit set). Our historical data, supported by our own pollinator observations, show that the orchids’ pollinator emerges after the start of orchid flowering, explaining selection for later flowering through fruit set. The difference in bee-emergence and orchid bloom seems, however, to not have changed in the last decades due to climate warming. We thus suggest that selection for later flowering is a consequence of low synchronization, which is however not caused by climate change, but has persisted for other reasons, such as slow adaptive response in the orchids, migration, or other, unmeasured selection against later flowering.

Our study contrasts with earlier work on the closely related *Ophrys sphegodes* and its pollinator bee, *Andrena nigroaenea*, from England, that documented the bee emerging significantly earlier than the flowering of the orchid (Robbirt et al., 2014). In addition, in this orchid-pollinator association, climate change seems to advance the flight time of the bee more strongly than the flowering time of the orchid, suggesting an increasing desynchronization between them in the future (Robbirt et al., 2014, Hutchings et al., 2018). For *Ophrys sphegodes* in England, Robbirt et al. (2011) calculated a 6d advance in flowering per 1°C temperature increase in spring. In Switzerland, the mean annual temperature has increased by about 2°C in the last 150 years (online data MeteoSwiss), and our observed advance in *Ophrys araneola* flowering is 12d, which perfectly matches the estimated advance in *O. sphegodes* flowering. For *A. nigroaenea*, Robbirt et al. (2014) showed an advance in emergence of up to 11.5d C^−1^, which is about double the value we document for *A. combinata*. Although our dataset with 52 bee-records is less extensive than the one used by Robbirt et al. (2014) which comprised 357 bee-records, the two bee species may well respond differently to increased temperatures. In bees, the response in emergence times to warmer winter temperature varies between species, with those that overwinter as adults tending to emerge earlier, whereas those that overwinter as larvae tending to emerge later (Frund et al., 2013). Whether the two species, *A. combinata* and *A. nigroaenea*, really respond differently to warmer spring temperature and what the reason for this may be, needs more investigation in the future.

The pattern of changed flowering and pollinator activity documented by Robbirt et al. (2014) suggest selection for earlier flowering in *Ophrys sphegodes* driven by pollination success, but perhaps counter-acted by frost damage. However, fruit set and frost damage was not measured in their study, and thus selection could not be quantified. Our data shows, in contrary, selection for later flowering, caused by early flowering individuals having a high likelihood of frost damage, and possibly a low chance of pollination. Our documentation of the pollinator bee in one of the populations, Birmenstorf, despite being anectodical, aligns with the peak record of this bee species in N-Switzerland (Figure 5) and thus supports the later occurrence of this bee species in relation to the flowering of the orchid. The observation of the pollinator bee in the population with the highest figures of fruit-set also supports *Andrena combinata* as the only (known) pollinator of *Ophrys araneola* (Schiestl and Vereecken, 2008), and the assumption that patterns of fruit set in our study populations are caused by natural pollination, rather than undocumented hand-pollination by orchid enthusiasts.

A puzzling question is why later flowering has not evolved in the orchids, despite the here documented strong selection for later flowering. This selection has likely persisted in this orchid throughout longer timescales, because the mismatch of bee-presence and orchid flowering seem unchanged in the past. It is possible that selection on later flowering through frost damage was weaker in the past, as frost damage may have been less severe with later flowering. However, because of global warming, spring frost was also stronger and more frequent in the past. Indeed, 60 years ago, there were 8 more frost days than today between March-May in Northern Switzerland (online data www.meteoschweiz.admin.ch), suggesting frost damage at that time has been at a similar level as now, even with later flowering.

The lack of evolutionary response to selection for later flowering may indicate low heritability but high environmentally-induced plasticity in this trait in *O. araneola*. Although heritability of flowering time has not yet been measured in this or any related orchid species, flowering time has been shown to be heritable and to evolve rapidly in other plant species, despite also being plastic (Franks et al., 2007, Anderson et al., 2012, Elzinga et al., 2007). Alternatively, selection that we did not account for may act against later flowering, such as early summer drought or herbivores, factors that may destroy later-ripening fruits. As another explanation for lack of evolutionary response, migration via seeds from earlier flowering, more southerly populations may prohibit or slow down adaptive evolution of later flowering.

In addition to documenting selection, our data of individually marked plants highlighted correlations between several kinds of traits of the same individuals among years. For example, flowering time was highly consistent throughout the years, and the same was true, but to a lesser degree, for plant size and number of flowers produced. As trait variability is caused by environmental-as well as genetic factors, consistent traits among years indicate either constant environmental conditions and/or genetic components of the traits. A constant environment is indeed suggested by our finding of correlated frost damage and fruit set in plants among years. Also, a trade-off between fruiting and subsequent flowering was indicated by our data by the fact that plants with fruits had a lower likelihood of flowering the next year again. Such a trade-off was already suggested by earlier studies in orchids, albeit in a population-, climate- and species-specific way (Sletvold and Ågren, 2015b, Sletvold and Ågren, 2015a, Calvo, 1993, Primack and Hall, 1990), and suggests that fruiting is costly in these orchids. Low fruit set has been found in other populations of *O. araneola* (Claessens and Kleynen, 2011) and may represent, in connection with long distance of pollen flow typically found in sexual mimics (Johnson and Schiestl, 2016), an adaptation for the production of fewer but high-quality seeds.

In the future it would be interesting to determine genetic variance and covariance of flowering time in this and other orchid species, e.g. by using genomics approaches, to find loci associated to flowering time and estimate the evolvability of this trait. This, together with figures of natural selection, will allow to build models on the speed of adaptive evolution, and thus to predict how fast populations can adapt in response to selection on flowering time. Such information can help to inform conservation measures such as assisted migration or hand pollination. Conservation should prioritize enabling and fostering of natural pollination by the pollinator bee, through conserving bee habitats and attempting to re-establish pollinator-bee populations where possible. As a last opportunity in case natural pollination fails, hand-pollination is an option to maintain reproduction in natural population. In case hand-pollination is done, our results suggest later-flowering individuals should be chosen for pollination, to potentially promote the evolution of a more favorable flowering time for natural fruit production.

## Acknowledgements

The authors would like to thank Michael Jutzi (www.infoflora.ch) as well as Christophe Praz and Julie Seemann-Ricard (www.infofauna.ch) for allowing us to use the historical records of *Ophrys araneola* and *Andrena combinata* form Switzerland for this paper. Franz Huber helped with field work and processing of temperature data. Salvatore Cozzolino and Donata Cafasso sequenced the DNA of pollinaria and a leaf of *Ophrys araneola*. Two anonymous reviewers helped to improve an earlier version of this manuscript. We also thank the nature conservation agencies of the Swiss cantons Aargau und Zürich for issuing the necessary permits to conduct this study. This study was funded by the University of Zürich.

---

**Supplementary Table S1.** Data of flowering time, frost damage, fruit set, number of flowers and plant size in the study populations and throughout the three years of the study. Mean values are followed by standard deviation.

| Year | Population<br>(number of plants<br>surveyed) | Mean day of<br>first flower | Percent<br>plants with<br>frost damage | Number of<br>plants with<br>fruits | Total number<br>of fruits | Mean<br>number of<br>fruits | Percent<br>individuals<br>with fruit-set | Mean<br>number of<br>flowers | Mean plant<br>size (cm) |
| --- | --- | --- | --- | --- | --- | --- | --- | --- | --- |
| 2021 | Erlinsbach (115) | 106.04±12.33 | 27 | 0 | 0 | 0.00±0.00 | 0 | 2.98±1.84 | 18.53±5.00 |
|  | Kloten (249) | 115.14±3.09 | 31 | 8 | 9 | 0.09±0.31 | 3 | 3.82±1.72 | 16.73±5.48 |
|  | Küttigen (123) | 107.11±9.31 | 75 | 1 | 1 | 0.04±0.21 | 1 | 5.44±1.61 | 24.15±6.05 |
|  | Birmenstorf (132) | 105.75±8.14 | 4 | 20 | 33 | 0.32±0.73 | 15 | 3.19±1.25 | 17.26±6.01 |
|  | Villigen (247) | 104.25±7.74 | 83 | 0 | 0 | 0.00±0.00 | 0 | 4.59±1.60 | 21.36±7.47 |
|  | Villnachern (77) | 90.59±6.81 | 80 | 3 | 3 | 0.04±0.19 | 4 | 5.34±1.70 | 16.95±5.91 |
| 2022 | Erlinsbach (109) | 102.60±12.01 | 27 | 0 | 0 | 0.00±0.00 | 0 | 3.95±1.95 | 11.11±3.76 |
|  | Kloten (284) | 107.88±4.14 | 1 | 5 | 6 | 0.03±0.19 | 2 | 4.53±1.81 | 18.80±6.24 |
|  | Küttigen (129) | 100.70±6.96 | 84 | 1 | 1 | 0.05±0.22 | 1 | 4.95±1.80 | 21.62±7.76 |
|  | Birmenstorf (132) | 101.26±7.59 | 12 | 13 | 20 | 0.29±0.67 | 10 | 3.57±1.50 | 17.64±5.08 |
|  | Villigen (274) | 99.25±7.17 | 41 | 2 | 2 | 0.02±0.12 | 1 | 4.91±2.25 | 23.24±7.72 |
|  | Villnachern (77) | 84.18±6.87 | 47 | 0 | 0 | 0.00±0.00 | 0 | 5.93±1.90 | 13.10±5.51 |
| 2023 | Erlinsbach (116) | 105.26±10.92 | 0 | 0 | 0 | 0.00±0.00 | 0 | 4.97±2.49 | 24.40±7.78 |
|  | Kloten (84) | 106.77±8.43 | 15 | 4 | 6 | 0.11±0.42 | 5 | 6.20±2.28 | 20.87±11.22 |
|  | Küttigen (165) | 101.25±6.17 | 13 | 4 | 4 | 0.04±0.20 | 2 | 5.38±2.55 | 26.56±8.00 |
|  | Birmenstorf (128) | 100.66±6.29 | 0 | 16 | 21 | 0.19±0.50 | 13 | 3.46±2.98 | 26.99±7.44 |
|  | Villigen (274) | 102.64±6.51 | 14 | 2 | 2 | 0.01±0.11 | 1 | 5.55±2.48 | 27.59±10.13 |
|  | Villnachern (77) | 92.20±5.51 | 47 | 4 | 4 | 0.06±0.23 | 5 | 6.30±1.55 | 24.68±8.10 |
| Overall mean ± s.d. |  | 101.90±7.11 | 33.38±29.87 | 4.61±5.93 | 6.22±9.19 | 0.07±0.10 | 3.43±4.6 | 4.73±1.03 | 20.64±4.72 |

**Figure S1.**
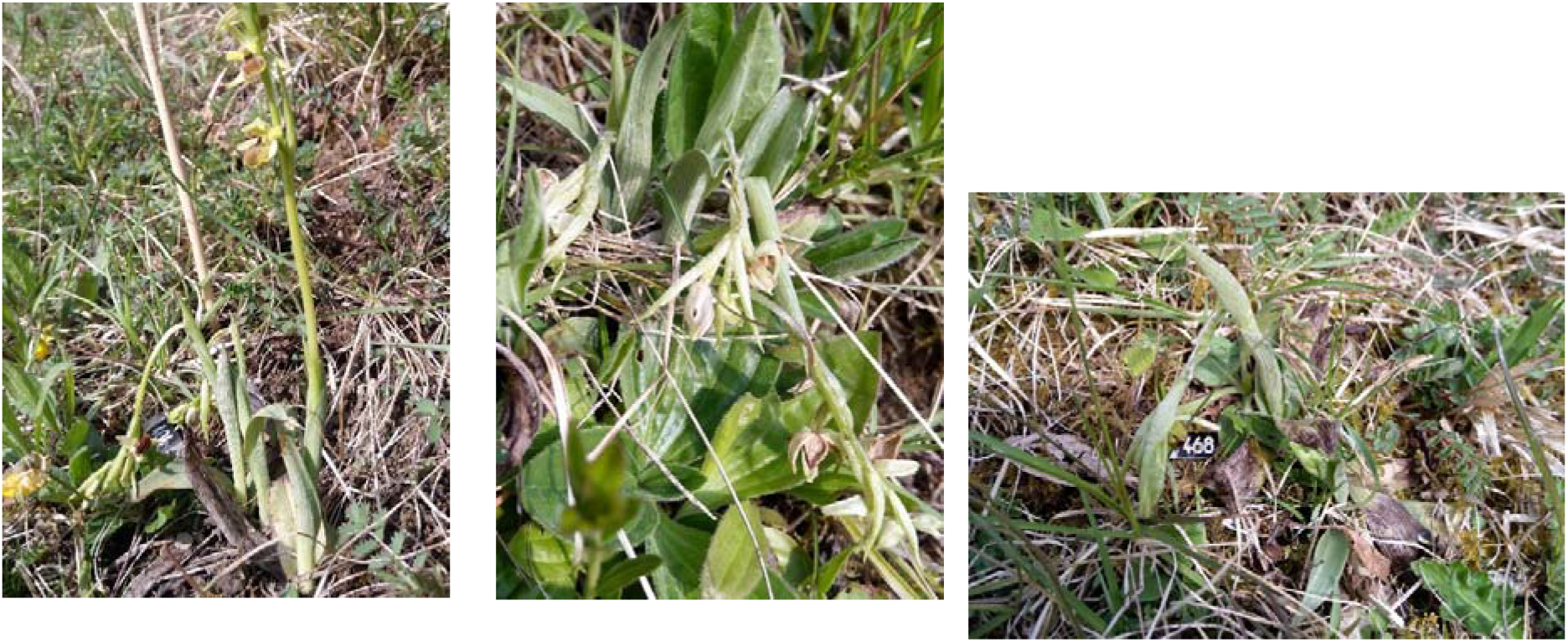
Pictures of frost-damaged plants in the population Villigen. In the right hand side picture, a numbering plate is visible.

**Figure S2.**
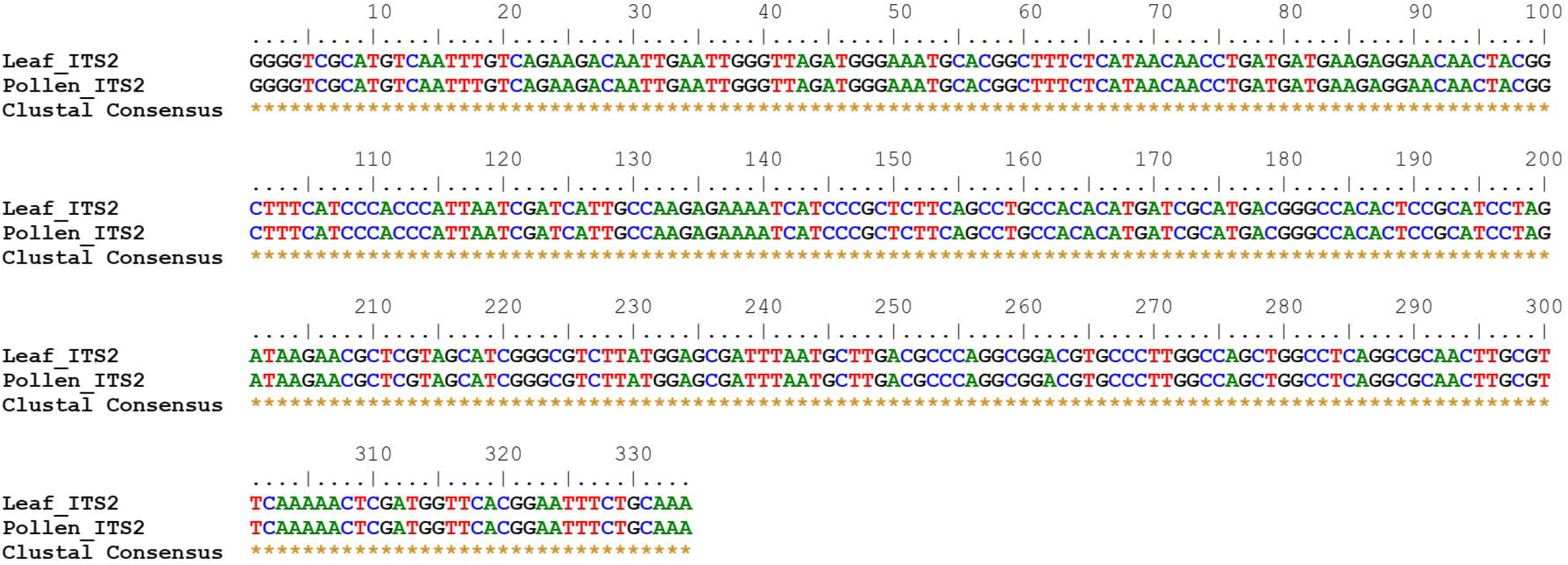
ITS2 sequences of a pollinarium carried by an *A. combinata* male caught in Birmenstorf, and a leaf sample collected from an *O. araneola* plant in the population.

